# Cholesterol-Mediated Modulation of Collecting Lymphatic Vessel Contractility: Exploring Cholesterol Depletion as a Therapeutic Alternative to Improve Lymphatic Function in Hypercholesterolemia

**DOI:** 10.64898/2026.08.04.742795

**Authors:** Keith Keane, Jorge Castorena-Gonzalez

## Abstract

Globally, hypercholesterolemia affects over 20% of the population; and while many studies have examined its impact on cardiovascular health, little is known about its effects on the lymphatic system. In mice, hypercholesterolemia has been linked to multiple aspects of lymphatic dysfunction; and a recent study demonstrated that cholesterol depletion by cyclodextrins promoted lymphatic vessel regeneration and restored lymphatic drainage in mouse models of lymphedema. Collecting lymphatic vessels rely on the spontaneous and highly entrained contractions of lymphatic muscle cells (LMCs) and competent unidirectional on-way valves to propel lymph forward. Critical to lymphatic pacemaking and contractility is the proper functioning of ion channels, which are known to be modulated by the cholesterol content in the plasma membrane. Therefore, we sought to understand the role cholesterol plays in regulating lymphatic contractility.

The effects of cholesterol depletion by the cyclodextrins MβCD and HPβCD were assessed in cannulated and pressurized inguinal-axillary collecting lymphatic vessels (CLVs) from C57BL6/J (WT) mice. Noteworthy, studies have shown that HPβCD is safe for human use, and in fact, it is commonly used as a drug excipient. Acute treatment with both cyclodextrins significantly increased the pumping capacity of CLVs, as demonstrated by the increased contraction amplitudes by ∼50±12% and calculated fluid volume displacement by each contraction by ∼35±11%. Calcium imaging demonstrated that HPβCD increased the amplitude and duration of the large Ca_v_1.2-mediated calcium events (termed calcium flashes. In contrast, cholesterol supplementation by incubation with BODIPY-cholesterol, which presumably incorporates cholesterol into the cell membrane, significantly impaired the contractile activity of CLVs compared to controls by decreasing contraction amplitude (control: 42±2 µm versus BODIPY-cholesterol: 20±7µm) and calculated fluid volume displacement (control: 9.2±3.9nL versus BODIPY cholesterol: 3.3±1.2nL) which were significantly restored with subsequent cholesterol depletion using HPβCD (amplitude: 36±11µm, volume displacement: 5.5±2.4nL). Similarly, treatment with HPβCD significantly improved the contractile capacity of dysfunctional CLVs isolated from hypercholesterolemic ApoEKO mice.

In conclusion, changes to cell membrane cholesterol content acutely and significantly altered CLV contractility with depletion improving contractility associated with recruitment of voltage-gated Ca_v_1.2 channels in lymphatic muscle cells (LMCs). Future studies from our lab will determine whether pharmacological depletion of membrane cholesterol can be therapeutic strategy to improve and/or restore lymphatic contractile function in secondary lymphedema, including obesity/hypercholesterolemia-induced and cancer-related lymphedemas.

## INTRODUCTION

The lymphatic system is integral to the trafficking of activated immune cells to local lymph nodes, the maintenance of tissue fluid homeostasis, and the absorption of macromolecules, particularly lipid nutrients in the gut and within other tissues^1–3^. Importantly, this system plays a key role in cholesterol homeostasis by trafficking nutrient cholesterol in the form of chylomicrons from the gut to the circulatory system as well as in reverse cholesterol transport where they remove cholesterol-loaded high-density lipoproteins (HDL) from our tissues and returns them to the bloodstream for excretion through the liver^4^. Overall, this system and the vessels that comprise it are key components in the proper management of cholesterol in health and disease; however, few studies have investigated the impact that abnormal accumulation of cholesterol within and surrounding the lymphatic vasculature and/or high lymph cholesterol have on lymphatic function, nor the impact of impaired lymphatic function on cholesterol homeostasis.

At present, over 20% of people worldwide have hypercholesterolemia^5^ often in association with obesity and metabolic syndrome. In the United States, about 10% of the adults over the age of 20 years have serum cholesterol levels higher than 240 mg/dL (6.2 mmol/L), the diagnostic threshold for hypercholesterolemia^6^; however, only half are actively using medication to manage it^7^. The relationship between serum cholesterol levels and vascular health has been under exploration since the early 1900s, it has since been linked to the development of atherosclerosis, coronary artery disease, but the exploration of its role in lymphatic dysfunction has just recently started^8–12^. In mouse models of hypercholesterolemia, immune cell trafficking and fluid transport are impaired and associated with a decrease in LMC coverage of CLVs while their contractions are characterized by a decrease in their contractile amplitude, strength, permeability, and valve patency of those vessels^13,14^, indicating a link between high cholesterol levels and lymphatic dysfunction.

Conversely, an impaired lymphatic system leads to the development of lymphedema, a condition defined by the buildup of protein-rich fluid within tissues that cannot be drained. This disease impacts not only the body’s ability drain fluid, but its ability to adequately respond to infection and inflammation due to impaired immune cell trafficking^15,16^ and the absorption and trafficking of lipid nutrients^17^ leading to lipid dysregulation^14,18^. It is estimated that primary lymphedema (genetic in origin) affects approximately 1:100,000 people while secondary lymphedema (due to lymphatic injury) affects 1:1,000; however, it is thought that these numbers may be underestimated^19^. In mice, a compromised lymphatic system is associated with adult-onset obesity^20^ and further compounds the accumulation of serum cholesterol in hypercholesterolemic mice^21^. In humans, large fat deposits are found around the compromised vessels of patients with lymphedema, a disease defined by inadequate lymphatic function^22^. Furthermore, while serum cholesterol levels in lymphedema patients are not diagnostically elevated, they are significantly increased compared to healthy controls; and tissue levels of cholesterol within the lymphedematous skin are significantly increased, and the restoration of lymph drainage ameliorates this accumulation^23^. Furthermore, treating mice with lymphedema with a cholesterol-scavenging molecule known as cyclodextrin, the lymphedema can be elevated.

Together, these studies highlight the close link between lipid dysregulation and lymphatic dysfunction; however, the mechanisms by which hypercholesterolemia impairs the contractility and pumping capacity of the CLVs remain unknown. In the vascular system, a diet high in cholesterol leads to the accumulation of cholesterol in the cell membrane of aortic smooth muscle cells^24,25^, decreases the expression of contractile proteins, and induces their differentiation to non-contractile foam cells^26,27^. This alteration in smooth muscle phenotype as well as the loss of LMCs in hypercholesterolemic mice^14^ implicates LMCs as a potential source of impairment of CLV contractile function but how this excess cholesterol is impacting these cells requires exploration.

Within cells, 90% of unesterified cholesterol is found within the plasma membrane where it directly contributes to membrane fluidity and the formation of cell signaling platforms known as lipid rafts^28^ and caveolae^29^; however, its function goes beyond these physical and structural components. Cholesterol is required for the proper functioning of a variety of proteins via direct binding, or through the association with cholesterol recognition motifs^30^. In fact, the functionality of many of the ion channels required for CLV contractility and pacemaking are altered by changes to membrane cholesterol including the voltage-gated calcium channel Ca_v_1.2, the voltage-gated calcium-activated chloride channel, anoctamin-1 (ANO1), TRP channels, BK channels, and hERG channels^31–33^.

Within the CLVs, Ca_v_1.2 is integral to the initiation of contractions as calcium influx is required to activate the smooth muscle contractile apparatus^34–36^ and the activity of Ca_v_1.2 has been demonstrated to be altered with cholesterol depletion and supplementation in a variety of different cells^37–41^. Specifically, cholesterol depletion increases calcium current density but also induces a positive shift in the current, indicating a larger influx of calcium ions but requiring a more positive threshold for activation^42^. On the other hand, cholesterol supplementation impairs or inhibits calcium currents^39,41^.

Previous studies have demonstrated that hypercholesterolemia has been linked to CLV contractile dysfunction^13,14,23^, therefore, in this study, we aimed to characterize the role cholesterol plays in regulating CLV contractile function. We assessed the effects of cholesterol depletion on WT vessels using the cyclodextrins methyl-ß cyclodextrin (MßCD) or (2-hydroxypropyl)-β-cyclodextrin (HPßCD)^43,44^, as well as supplementing cholesterol in WT vessels using BODIPY-cholesterol^45,46^. We tested the hypothesis that excess levels of cholesterol incorporated into the cell membrane result in abnormal functioning of ion channels, including those critical for lymphatic contractility and pacemaking. This impairment leads to severe contractile dysregulation, that can be rescued by pharmacological depletion of cholesterol by cyclodextrins. To test this hypothesis, we characterized the *ex vivo* contractile function of CLVs from hypercholesterolemic ApoE KO mice that develop impaired lymphatic function^13,14,47^ or from WT mice supplemented with BODIPY-cholesterol under baseline conditions and following acute treatment with HPßCD ^43,44^. Importantly, cyclodextrins are common ingredients found in commercial drugs where they are used as excipients ^48,49^, and HPßCD is currently under investigation in clinical trials to treat Niemann-Pick Disease Type C1, and Alzheimer’s disease, while other diseases such as Parkinson’s disease have shown some success in preliminary studies (reviewed by Nicolaescu OE, et al., Pharmaceutics, 2025^50^) and recently was shown to improve lymphatic function in a mouse model of lymphedema^23^. These highlight the unexplored therapeutic potential of HPßCD to enhance and/or rescue lymphatic contractile function and pumping capacity of dysfunctional CLVs in hypercholesterolemia as well as secondary lymphedema associated with obesity and metabolic syndrome.

## MATERIALS AND METHODS

### Animals

C57BL/6J (WT, Strain No.: 000664), ApoE^−/-^ (B6.129P2-Apoetm1Unc/J, Strain No.: 002052), Myh11-iCreERT2 (B6.FVB(Cg)-Tp(X)1SoffTg(Myh11-icre/ERT2)1Soff/ZjngJ, Strain No.: 036935), and Salsa6f (B6(129S4)-Gt(ROSA)26Sortm1.1(CAG-tdTomato/GCaMP6f)Mdcah/J, Strain No.: 031968) mice were purchased from The Jackson Laboratory. Cx45^fx/fx^ mice were obtained from Klaus Willecke, University of Bonn, Germany. Myh11-iCreER^T2^;Cx45^fx/fx^ and Myh11-iCreER^T2^;Salsa6f mice were generated through in-house breeding by crossing either Cx45^fx/fx^ or Salsa6f mice with Myh11-iCreERT2. Cre-recombination in these mice was induced by intraperitoneal injection of 1 mg/day of tamoxifen for 5 days. Tamoxifen (Cat. No.: T5648-1G; Sigma) injections were prepared at a concentration of 10 mg/mL in sunflower seed oil (Cat. No.: S5007-250ML; Sigma) with 10% v/v ethyl alcohol (Cat. No.: E7023-500ML; Sigma). Mice were housed in groups of maximum 5 mice per cage under a 12-hour light/dark cycle. Room temperature was maintained at 22–25°C with food and water available *ad libitum*. All mice in this study received a control, regular mouse chow (PicoLab RodentDiet 20 Cat. No. 355043) and used between 3 and 6 months of age. Mice were anesthetized with isoflurane and euthanized by overdose with isoflurane followed by cervical dislocation.

### Solutions and Chemicals

During microdissection and cannulation of lymphatic segments, a *Krebs-BSA* buffer was utilized. It contained: 146.9 mM NaCl (Cat. No.: 746398, Sigma) 4.7 mM KCl (Cat. No.: 746436, Sigma) 2 mM CaCl_2_ 2H_2_O (Cat. No.: C5080, Sigma), 1.2 mM MgSO_4_ (Cat. No.: 793612, Sigma), 1.2mM NaH_2_PO_4_ H_2_O (Cat. No.: 71507), 3 mM NaHCO_3_, 1.5mM Na-HEPES (Cat. No.: H8651, Sigma), 5 mM D-glucose (Cat. No.: RDD016, Sigma), and 0.5% BSA (Cat. No.: A3311, Sigma) at pH = 7.4). During pressure myography experiments, lymphatic segments were constantly superfused with a *Krebs* buffer without BSA. For imaging of BODIPY-Cholesterol treated vessels, cannulated and pressurized lymphatic segments were perfused with a Ca^2+^-free Krebs buffer where 3 mM EDTA replaced calcium (Cat. No.: E4884, Sigma). For acute cholesterol supplementation, lymphatic vessels were incubated with 10 µM BODIPY-Cholesterol (Cat. No.: 24618; Cayman Chemical) in Krebs buffer. For cholesterol depletion, we used the cyclodextrins MßCD (Cat. No.: HY101461; MedChem Express) and HPßCD (Cat. No.: HY-101103; MedChem Express) dissolved in Krebs buffer. Nicardipine (Cat. No.: N7510, Sigma) were prepared as a 20 mM stock solution in dimethyl sulfoxide (DMSO, Cat. No.: D2438-10mL; Sigma) and further diluted in Krebs buffer to reach a final working concentration of 1 µM.

### Pressure Myography

Inguinal axillary collecting lymphatic vessels (i.e., iaCLVs) were used for all the experimental protocols included in this study. iaCLVs were isolated as previously described^35,51–53^. Briefly, after euthanasia, with a mouse in the prone position, a skin cut was initiated near the base of the tail and extended cephalad to the scapula. A small cut was made to allow the tissue to be stretched and pinned on a dissection board. The area was flooded with a Krebs-BSA buffer to keep the area moist. The iaCLVs run between two fat pads (one in the inguinal region and one in the axilla) and interconnect the inguinal and axillary lymph nodes. Starting at the inguinal lymph node, the lymphatic vessel and surrounding tissue between the inguinal and axillary lymph nodes were excised from the animal and pinned down onto a Sylgard-coated dissection chamber filled with Krebs-BSA buffer. Fine-tipped scissors were used to isolate the lymphatic vessel from the surrounding tissue and vessels were stored in Krebs-BSA buffer at room temperature until cannulation for pressure myography experiments. For cannulation and functional assessment, iaCLV segments were transferred to a custom designed 2.5 mL pressure myography chamber filled with Krebs-BSA buffer. Vessel segments were cannulated and pressurized to 3 cmH2O at 37°C under no intraluminal flow conditions, using two glass micropipettes (input ∼75 µm outer diameter; output ∼90 µm diameter outer diameter), and allowed to equilibrate for 30 minutes under constant perfusion of BSA-free Krebs buffer (180 µL/min). An OB1 MK3 microfluidic flow control system (Elveflow, Paris) was utilized for precise control of the intraluminal pressure. Brightfield videos of vessel contractions were recorded at 20 frames-per-second and analyzed for lymphatic contractile parameters.

### Intracellular Calcium Imaging

Characterization of lymphatic muscle cell (LMC)-specific intracellular calcium dynamics was completed using collecting lymphatics from Myh11-iCreER^T2^;Salsa6f mice. Imaging was performed using a high-speed, high-resolution spinning disk confocal microscope (i.e., ANDOR Dragonfly 202, Oxford Instruments). Videos of changes in fluorescence were collected at 10-20 fps using a Leica HC FLUOTAR L 25x/0.95 objective and an iXon Life 888 EMCCD Camera. 2-dimentional maps (i.e., space-time maps – STM) were generated^53^, representing the time-dependent fluorescence intensity from a line-scan analysis at every position along the vessel were generated. Intracellular calcium events were associated with an increase in fluorescence intensity. The changes in fluorescence were measured over time (via line-scan analysis) and then represented as STMs with fluorescence intensity encoded in a grayscale (with white/light-gray colors associated with maximal fluorescence intensity over a dark background).

### BODIPY-Cholesterol Depletion

Isolated iaCLVs were mounted on glass pipettes and a 5 µM solution of BODIPY-Cholesterol (Cayman Chemical, Cat. No.: 24618) in Krebs-BSA buffer was applied to the lumen and incubated in the BODIPY-cholesterol solution for 1 hour at 37°C. Vessels were flushed with Krebs BSA buffer, transferred to a second chamber, mounted on glass pipettes containing calcium-free Krebs buffer. Vessels were equilibrated at 37°C in Krebs-BSA before being transferred to calcium-free Krebs buffer. After contractions ceased, vessels were imaged at high resolution using an inverted spinning disk confocal microscope ANDOR Dragonfly 202 (Oxford Instruments), a 488 nm laser, and a Zyla PLUS 4.2 megapixel sCMOS camera using a Leica HC FLUOTAR L 25x/0.95. Images were taken before cholesterol depletion with 5 mM HPßCD or vehicle control, and every 15 minutes after treatment. The mean fluorescent intensity (MFI) minus the background MFI was assessed using ImageJ and compared to time zero to determine the percent change from baseline.

### Single-cell RNA sequencing (scRNAseq)

Data for scRNAseq was previously generated and described by our group^54^. Data analysis was performed using Seurat (4.3.0) in R studio (RStudio 2025.09.1+401).

### Statistical Analyses

Results were analyzed with GraphPad Prism, Version 10.6.1. Wilcoxin signed rank test, 2-way ANOVA corrected with Tukey test for multiple comparisons, and 1-way ANOVA test that included a correction for multiple comparisons using Dunnett test and a Geisser-Greenhouse correction as no equal variances were assumed (i.e., sphericity was not assumed) were used to determine differences in contractile function parameters, with statistical significance set at *p*<0.05. Statistical tests were specified in the captions of each figure legend. For pressure myography experiments, the number N of experiments corresponds to each individually tested collecting lymphatic segment while n corresponds to the number of animals tested.

## RESULTS

### Cholesterol depletion alters lymphatic contractile function in a concentration- and time-dependent manner

To understand how changes to cell membrane cholesterol alter the function of the CLVs, we first assessed the effects of cholesterol depletion using the cyclodextrins MßCD and HPßCD. Cyclodextrins are known to deplete cholesterol in a concentration and time dependent manner, with lower concentrations and shorter time periods depleting cholesterol in a more selective manner ^55^. Therefore, we exposed iaCLVs to gradually increasing concentrations of cyclodextrins in a range from 0 to 10mM. A single acute dose of each cyclodextrin concentration was given every 5 minutes (with contractions being analyzed in the last 2 minutes), while the preparation was maintained under constant superfusion with fresh Krebs buffer. Compared to control vessels (Fig. 1A), cyclodextrins MßCD (Fig. 1B) and HPßCD (Fig. 1C) significantly increased contraction amplitude at concentrations greater than 1 mM for MßCD and 5 mM for HPßCD (Fig. 1D). Conversely, higher concentrations of cyclodextrins resulted in a significant decrease in contraction frequency, with MßCD and HPßCD significantly decreasing frequency at 10 mM and 5 mM respectively (Fig. 1E). In a recent study, we introduced a new calculated parameter that aims to estimate the volume of fluid being displaced by each contraction ^56^. The calculated volume displaced was significantly increased in the MßCD group at concentrations higher than 3 mM (Fig. 1F).

**Figure 1:**
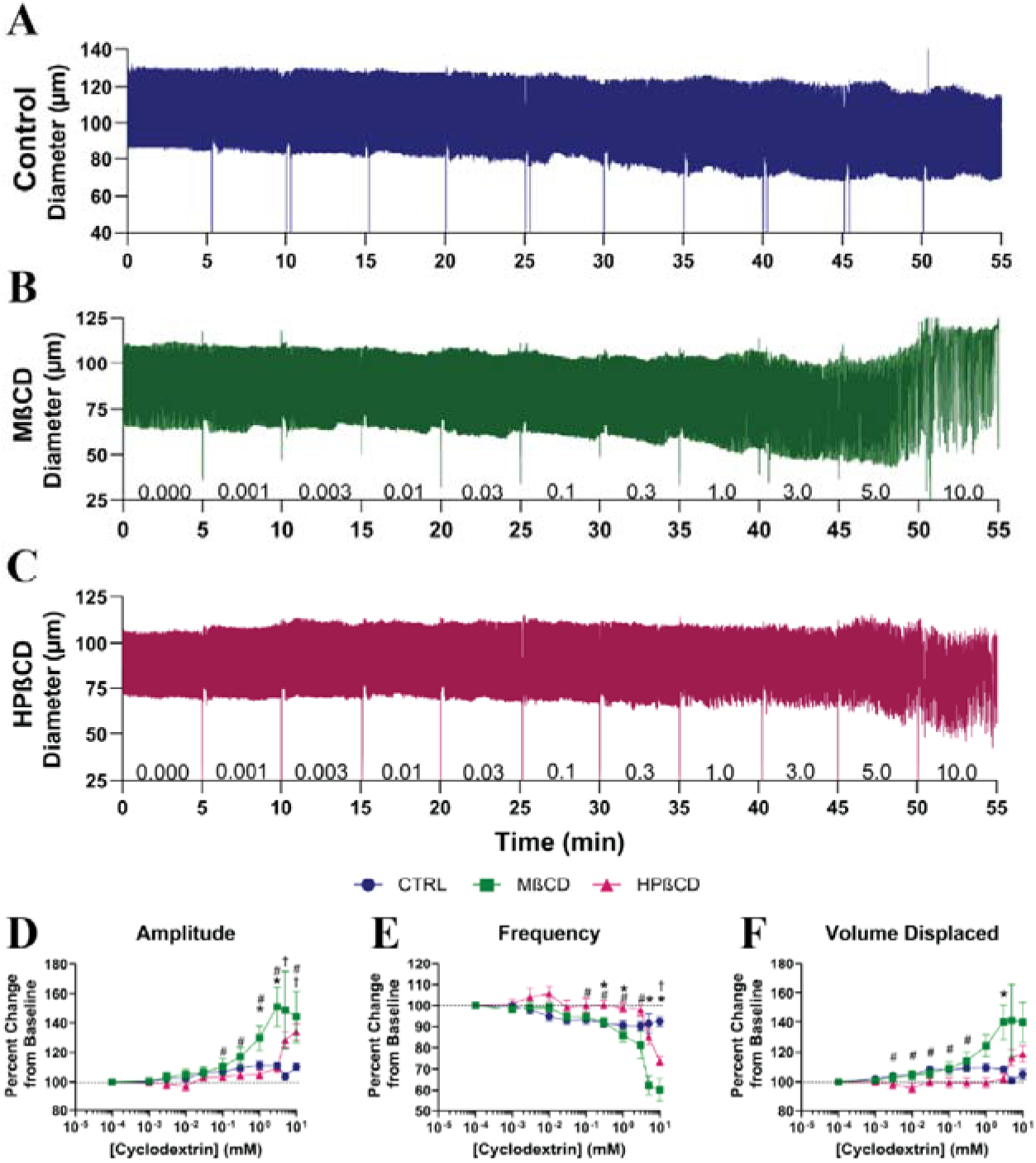
Concentration response of iaCLVs to cyclodextrins. Representative concentration response diameter traces of iaCLVs treated with A) vehicle control, B) MßCD, and C) HPßCD. Vessels were assessed for changes in D) amplitude, E) frequency, F) volume displaced. Data is represented as the mean ±SD of n=18 animals, Control: N=10 vessels, MßCD: N=10 vessels, HPßCD: N=8 vessels per group. A two-way ANOVA using Dunnett’s multiple comparisons test was performed. Statistical significance was set at *p*< 0.05 and compared to the baseline for each vessel and are identified as follows: control (#), MßCD (*), HPßCD (†).

The depletion of membrane cholesterol changes the organization and fluidity of the plasma membrane and alters the localization and movement of proteins embedded within it ^8,57–59^. While some changes in contractile function were observed even after short 5-minute stimulations (Fig. 1), these processes and their potential chronic effects are likely better assessed with longer observations. Therefore, we assessed the changes to lymphatic contractile function after a single washout dose of either vehicle control, 1 mM MßCD, or 5 mM HPßCD (i.e., cyclodextrin concentrations at which contraction amplitude was maximal in Fig. 1), while contractions were monitored for 60 minutes (Fig. 2). Within the first 10 minutes after treatment, both MßCD and HPßCD demonstrated a significant increase in contraction amplitude (Fig. 2B-D). Lymphatic vessels were constantly superfused with a fresh Krebs buffer (at 180μL/min), which would imply that most of the drug (i.e., cholesterol depletion agent) that was initially incorporated into the 2.5mL observation chamber would have been washed out after ∼14 minutes. Interestingly, the significant increase in contraction amplitude was maintained compared to that of baseline throughout the 60-minute recording, while contractile function in vehicle-treated vessels was unchanged (Fig. 2A-D). Vessel end diastolic diameter was unaltered after cholesterol depletion (Fig. 2E), which implied no changes in myogenic tone and indicated that the increase in amplitude after MßCD and HPßCD treatment was due to the significant decrease in end systolic diameter (Fig. 2F). These results also pointed to stronger lymphatic contractile capacity after cholesterol depletion.

**Figure 2:**
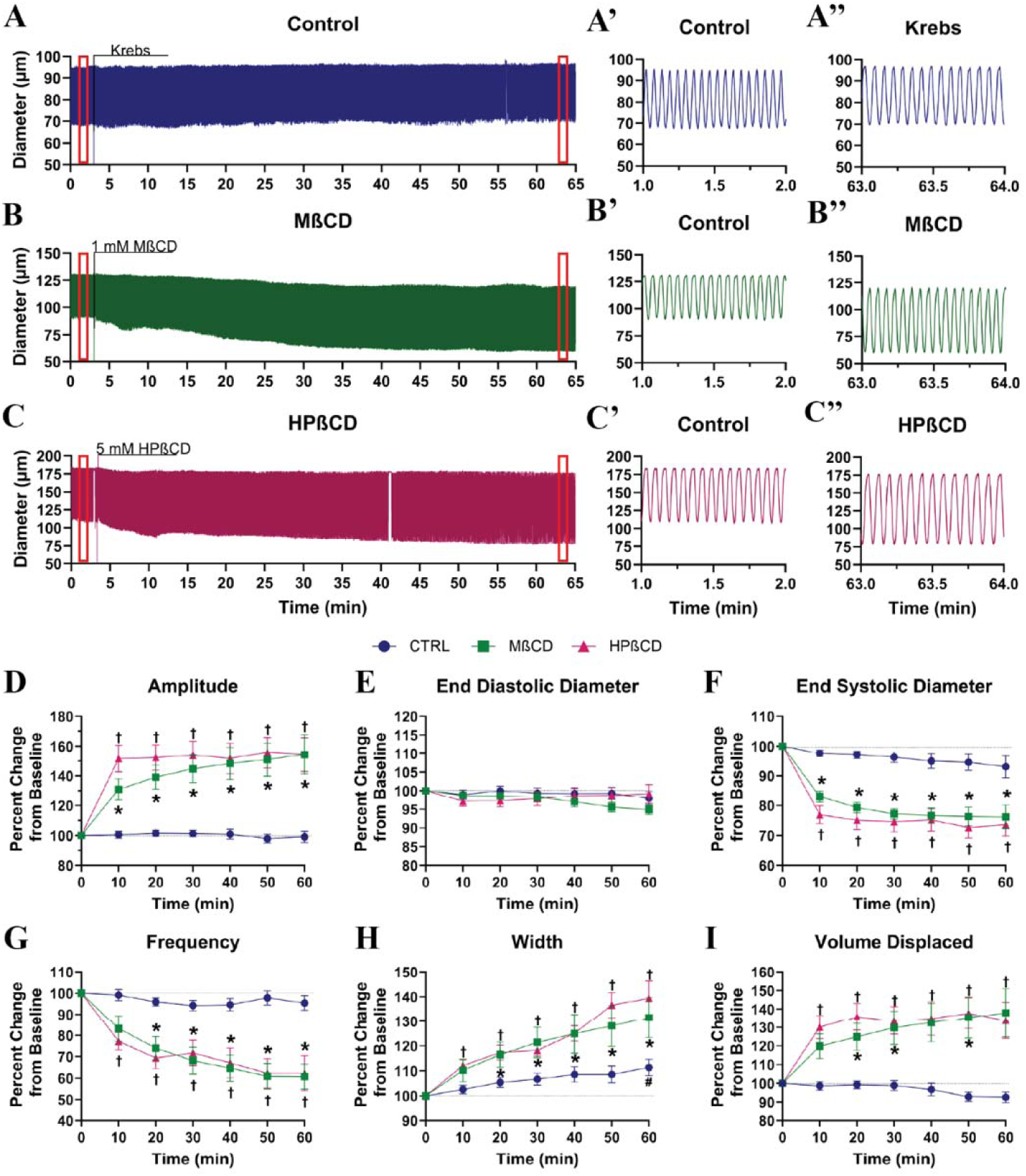
Cyclodextrins increase contraction amplitude but decrease frequency. Representative 60-minute diameter tracking of an iaCLV exposed to A) vehicle control, B) MßCD, and C) HPßCD. Red boxes highlight 1-minute diameter tracking at A’,B’,C’) baseline control and A”,B”,C”) after 60 minutes of vehicle or cyclodextrin treatment. Vessel traces were assessed for changes over time in D) amplitude, E) end diastolic diameter, F) end systolic diameter, G) Frequency, H) contraction width, and I) volume displaced. Data are expressed as ±SEM of n= 6-8 animals, Control: N=10 vessels, MßCD: N=10 vessels, HPßCD: N=10 vessels. A two-way ANOVA with Dunnett’s multiple comparisons test performed. Statistical significance was set at *p<* 0.05 and compared to the baseline for each vessel and are identified as follows: control (#), MßCD (*), HPßCD (†).

Control vessels showed no significant change in contraction frequency compared to baseline over 60 minutes; however, cholesterol depletion with either MßCD or HPßCD resulted in decreased contractile frequency, and this was maintained for the duration of the observed time (Fig. 2G). The decrease in frequency in the cyclodextrin-treated vessels was associated with a significant increase in the width of each contraction (Fig. 2H), indicating an increase in the length of the contraction cycle compared to vehicle-treated control vessels, rather than just a decrease in the number of contractions per minute. Changes to contractile efficiency were assessed by calculating the change in displaced volume per contraction, with an increase in displaced volume correlating with improved contractile efficiency. Both MßCD and HPßCD significantly increased the volume displaced for each contraction compared to vehicle-treated vessels (Fig. 2I).

### Cyclodextrins decrease membrane cholesterol in live collecting lymphatics

Next, we wanted to confirm that exposing live lymphatic vessels to cyclodextrins did in fact result in depletion of membrane cholesterol from the different cells that make up the lymphatic wall. Traditionally, cholesterol depletion is confirmed using Filipin III, a naturally fluorescent bacterial-derived toxin which binds to cholesterol; however, due to its cellular toxicity this approach is not suitable for live tissues. Therefore, we decided to incubate iaCLVs with 5 µM BODIPY-cholesterol, a fluorescently labelled cholesterol which incorporates into the plasma membrane of living cells, allowing for a direct quantitative assessment of membrane cholesterol over time in a live vessel ^45,46^. Vessels with incorporated BODIPY-cholesterol were cannulated, pressurized and allowed to equilibrate at 37°C for 30 minutes in Krebs buffer. Cholesterol content was then assessed before and after treatment with cyclodextrins using high-speed and high-resolution fluorescence confocal microscopy. Prior to imaging, the Krebs buffer in the observation chamber was replaced with a 37°C calcium-free Krebs to inhibit contractions and maintain the preparation stable, allowing for imaging of the live vessel for the length of the experiment, i.e., 1 hour. While our results demonstrated that lower concentrations of MßCD (i.e., 1mM) were required to induced comparable effects than those induced by higher concentrations of HPßCD (i.e., 5 mM), only HPßCD is considered safe for human consumption, approved by the FDA, and its use is currently being assessed in clinical trials ^50,60^. Therefore, we decided to focus on HPßCD for the remainder of this study.

Vessels treated with a single dose of 5 mM HPßCD displayed a significantly greater loss in mean fluorescent intensity (MFI) of BODIPY-cholesterol over time compared to vehicle control-treated vessels (Fig. 3A-C). Importantly, most of the loss in MFI occurred within the first 15 minutes (Fig. 3C). This is consistent with our functional experiments, where the greater change in contractile parameters took place within the first 10-20 minutes (Fig. 2), and with the fact that, under constant superfusion with a fresh buffer at a rate of 180 μL/min, the estimated bath turned over and drug washed out would be ∼14 min. While the loss of MFI in the vehicle-treated mice was significant compared to baseline, the amount of signal (i.e., incorporated BODIPY-cholesterol) lost between each timepoint was comparable and not significant (Fig. 3D), suggesting that this loss may have been associated with either the processing of the BODIPY-cholesterol by the cells ^61^ or photobleaching, although BODIPY dyes are known for their high photostability and strong resistance to photobleaching.

**Figure 3:**
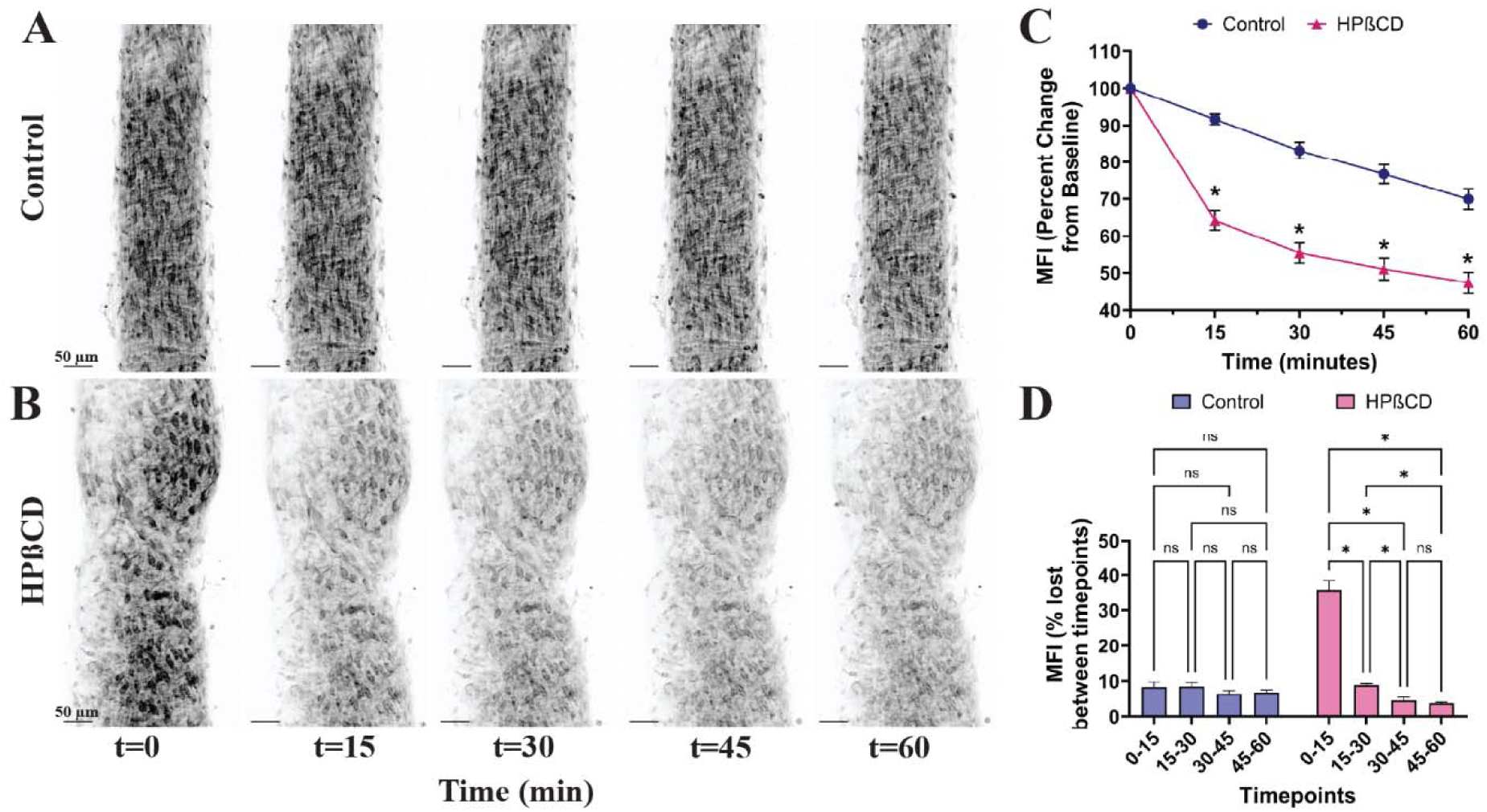
HPßCD decreases membrane cholesterol in iaCLVs. Representative confocal images of iaCLVs incubated with 5µM Bodipy-cholesterol and treated with A) control or B) 5 mM HPßCD washout over one hour. C) Changes to baseline MFI were compared between vehicle and HPßCD treatment for each timepoint and D) the loss of MFI between timepoints of each treatment. A two-way ANOVA using Tukey’s multiple comparisons test was performed. Data are represented as the mean ± SEM of N=6 animals, n=6 vessels per group. \**p*<0.05. Scale: 50 µm.

### The increase in the contractile capacity of collecting lymphatics induced by cyclodextrins is associated with increased intracellular calcium mediated by Ca_v_1.2 channels

CLV contractions are initiated by action potentials that rapidly propagate along the LMC layer entraining the coordinated influx of calcium through Ca_v_1.2 channels^34–36,53^. Ca_v_1.2 channels have been reported to be sensitive to changes in membrane cholesterol, with cholesterol depletion increasing channel activity and recruitment in cardiac and coronary myocytes and auditory hair cells^39,40,42^ while cholesterol supplementation decreases calcium entry in mouse coronary myocytes and guinea pig gallbladder cells^39,62^.

Therefore, we next assessed the modulation of the intracellular calcium dynamics in LMCs by HPßCD using Myh11-iCreER^T2^;Salsa6f mice (which express the genetically encoded calcium indicator GCaMP6f and a tdTomato reporter) and high-speed and high-resolution fluorescence confocal microscopy. We first characterized the effects of HPßCD on the large Ca_v_1.2-dependent calcium events that drive lymphatic contractions termed *calcium flashes* ^35,53,63^. The rhythmic, spontaneous contractions of collecting lymphatics make imaging of intracellular calcium activity particularly complex. To avoid incorporating additional pharmacological agents to blunt the sample movements associated with these contractions, e.g., wortmannin, we decided to focus on the middle of the vessel (i.e., half point between the lower and upper sides of the wall), which allowed us to image a consistent vessel plane (Fig. 4A). The spontaneous calcium activity of iaCLVs was recorded for 1 minute under control conditions and for 2 minutes following a single stimulus with 5 mM HPßCD. We generated two-dimensional maps, i.e., space-time maps (STMs), from line-scan analysis of fluorescence intensity, as previously described^35,53,63,64^. These STMs allow for visual representation of the changes in fluorescent intensity, which are associated with changes in intracellular calcium. In these STMs, each white/light-gray vertical band represents a calcium flash (Fig. 4B). Consistent with our results in Fig. 2, treatment with HPßCD resulted in a significant increase in calcium flash amplitude (represented as F/F_0_) and width, and a significant decrease in calcium flash frequency (Fig. 4C-E). Further implicating HPßCD in the recruitment of Ca_v_1.2 channels.

**Figure 4:**
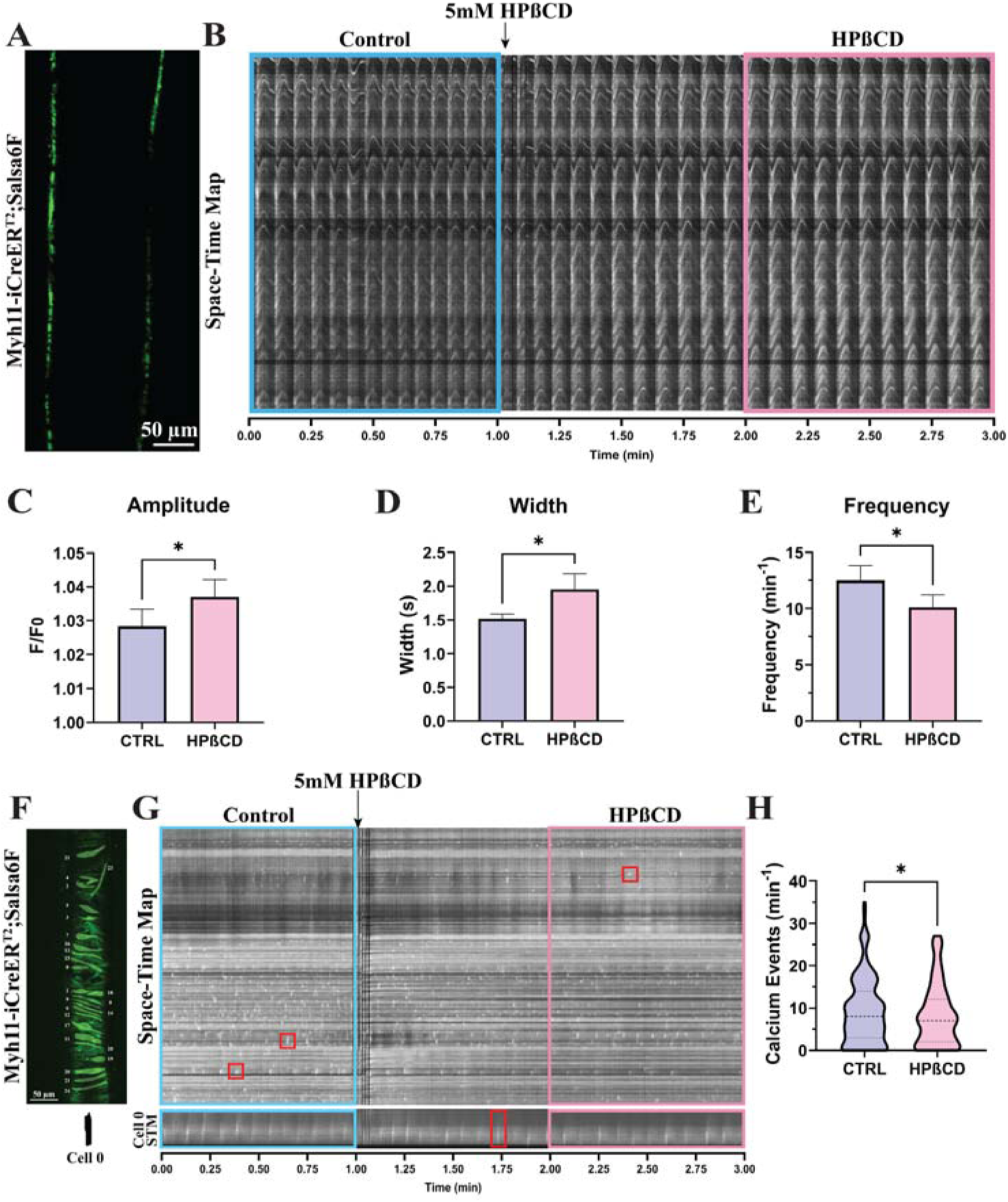
Cholesterol depletion alters LMC calcium dynamics. A) Representative IF confocal microscopy image of a Smmc^Cre.ERT2^;Salsa6f iaCLV. B) Representative 3-minute STM of an iaCLV treated with 5mM HPßCD at 1 minute. Blue and pink boxes represent assessed frames for C) amplitude, D) width, and E) frequency of calcium flashes. F) Representative image of Smmc^Cre.ERT2^;Salsa6f iaCLV with an overlay of individual LMC-specific masks (blue). G) Representative 3-minute STM of a 1 µM nicardipine-treated iaCLV treated with 5mM HPßCD at 1 minute with F’&G’) representative mask and STM of a single cell. Blue and pink boxes represent assessed frames for control and HPßCD respectively of assessed H) calcium events. Data are represented as the mean of ± SEM of A-E) N=8 vessels from n=3 animals and F-H) the mean of ± SEM of N= 9 vessels from n=3 animals, and I) N= 136 cells. Wilcoxon rank test was performed. * *p* < 0.05.

Next, we aimed to determine if HPßCD may also modulate intracellular calcium in LMC independently of Ca_v_1.2 channels. Therefore, we characterized the intracellular calcium activity in iaCLVs in the presence of the Ca_v_1.2 channel inhibitor nicardipine under control conditions (1 minute) and after a single stimulus with 5 mM HPßCD (2 minutes). Nicardipine (1 µM) inhibited all calcium flashes and associated contractile activity, allowing for the imaging of Ca_v_1.2-independent spontaneous calcium events on the lower wall face (Fig. 4F). A representative STM of the intracellular activity in the entire vessel is shown in Fig. 4G (upper panel), where the discrete, localized white/light-gray particle-like features are indicative of spontaneous calcium transient events (examples are shown in red boxes). Our initial impression was that exposure to HPßCD decreased the overall intracellular calcium activity. To confirm this, we generated STMs for individual cells and quantitatively assessed the frequency of intracellular events (Fig. 4G, bottom panel). In the representative example shown in Fig. 4, a total of 25 cells were analyzed (Fig. 4F). On average, cells displayed a total of 9.1±7.7 events/minute under control conditions, while the number of events was significantly reduced to 8.3±7.3 events/minute following treatment with HPßCD (Fig. 4H). These results point to cholesterol depletion by HPßCD as a modulator not only of Ca_v_1.2 channels, but also other channels that participate in the control of intracellular calcium, e.g., Ano1 and BK channels^63,65,66^.

### The pressure-dependent contractile function of collecting lymphatics is modulated by cholesterol

Transmural pressure and flow-mediated shear stress are among the key modulators of CLV contractility ^63,67–71^. As described earlier, the experiments included in this study were completed in the absence of intraluminal flow (i.e., both inflow and outflow pressures were equal), and therefore, in the absence of flow-mediated shear stress. We assessed the pressure-dependent response in contractile function of iaCLVs under control conditions and after cholesterol depletion with HPßCD. Pressure responses were assessed at 3, 0.5, 1.0, 2.0, 3.0, and 5.0 cmH_2_O for each treatment. For cholesterol depletion, vessels were given a single stimulus with 5 mM HPßCD while maintaining constant superfusion with fresh Krebs buffer. The pressure response after HPßCD treatment was completed 30 minutes after the drug was administered to allow for the drug to reach a maximum and stead effect in response to cholesterol depletion (based on the data in Fig. 2). Consistent with previous studies^35,53,63,68,69,72,73^, lymphatic vessels displayed an exquisite sensitivity to changes in intraluminal pressure (Fig. 5A). As pressure increased, so did frequency and end diastolic diameter. Contraction amplitude was maximum between pressure 1 and 2 cmH_2_O and myogenic responses were more evident when pressures increased to 3 and 5 cmH_2_O. These responses were similar following cholesterol depletion. Interestingly, contraction amplitude was increased by 25-50% at pressures ≥ 2 cmH_2_O (Fig. 5C), suggesting that cholesterol depletion with HPßCD could be a potential therapeutic approach to improve the pumping capacity of collecting lymphatics, especially when the lymphatic networks are overloaded, i.e., higher transmural pressures. End diastolic diameter was significantly decreased at most pressures, indicating an increase in myogenic tone (Fig. 5D). Given the increase in amplitude and decrease in end diastolic diameter, end systolic diameters were decreased at all pressures (Fig. 5E). Consistent with the contraction and calcium imaging data in previous sections, contraction frequency trended lower after cholesterol depletion and was significantly decreased at pressures 3 and 5 cmH_2_O, while contraction width was significantly increased at all pressures (Fig. 5F,G). Interestingly, the calculated fluid volume displaced per contraction was significantly decreased at low pressures (i.e., <1 cmH_2_O), but was significantly increased at higher pressures (i.e., >2 cmH_2_O, Fig. 5H).

**Figure 5:**
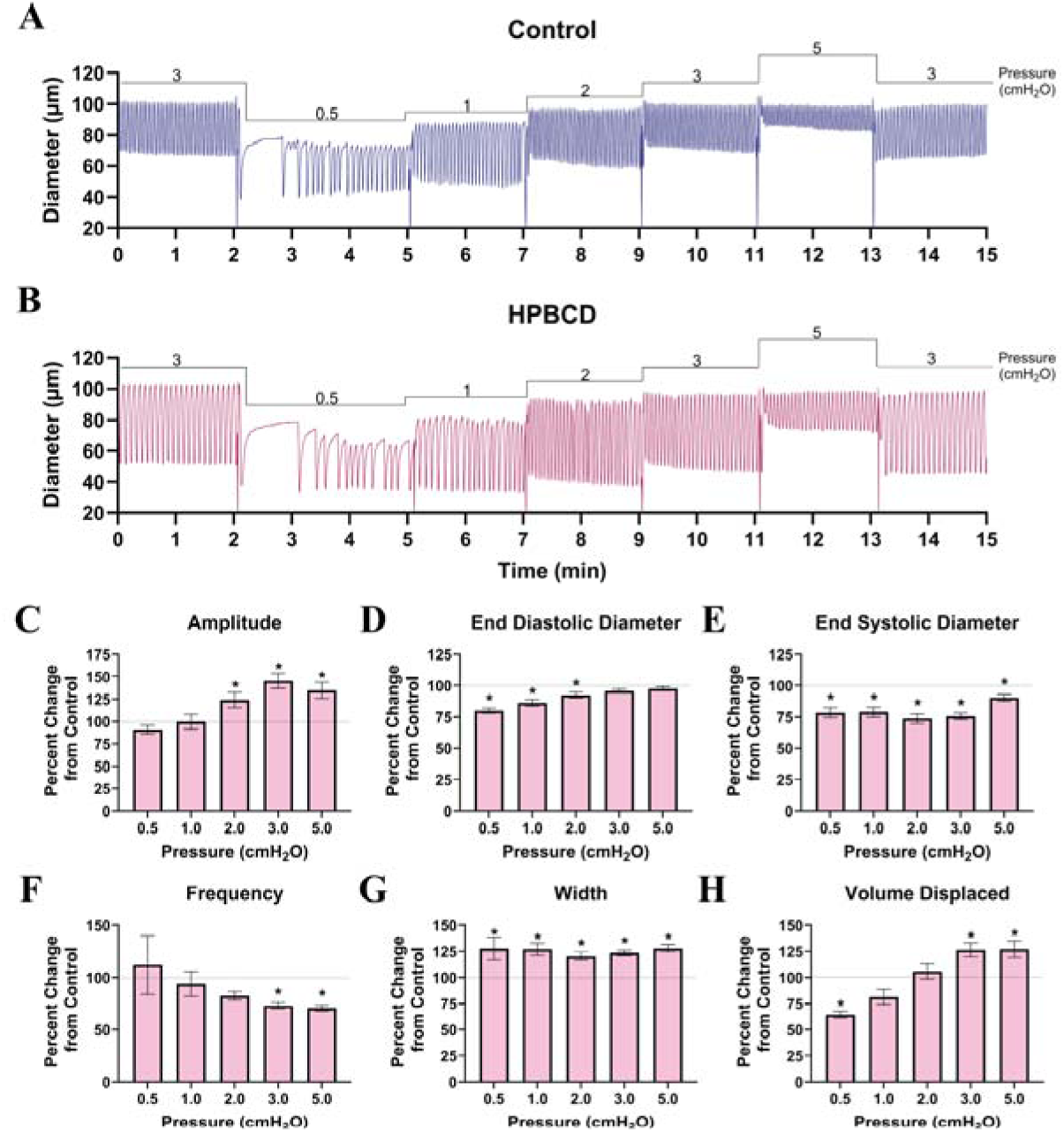
Cholesterol depletion alters iaCLV pressure response: Representative diameter tracking of an iaCLV exposed to increasing luminal pressure under no-flow conditions at A) control and 30 minutes after B) 5mM HPßCD treatment. Vessel traces were assessed for changes over time in C) amplitude, D) end diastolic diameter, E) end systolic diameter, F) Frequency, G) contraction width, and H) volume displaced per contraction. Data are expressed as ±SEM of n= 9 animals, N=10 vessels. A two-way ANOVA with Sidak’s multiple comparisons test was performed. Statistical significance (*) was set at *p<* 0.05 and compared to the baseline for each vessel.

### Changes in lymphatic contractile function induced by cyclodextrins are independent of LMC-to-LMC communication mediated by Connexin-45

Lymphatic contractions rely on the directional propagation of membrane depolarizations, i.e., pacemaking action potentials, along the lymphatic muscle cell (LMC) layer, which entrains the coordinated contraction of LMCs ^35,53,64,74^. We previously demonstrated that connexin 45 (Cx45) is the major connexin expressed in LMCs and mediates LMC-to-LMC coupling. Loss of Cx45 in LMCs resulted in very erratic, uncoordinated contraction^53,75^. Relevant to this study, some reports suggested that connexin gap junctions involved in intercellular coupling could be localized within cholesterol-dense lipid raft microdomains^76^. Therefore, to determine whether coupling between LMCs through Cx45 gap junctions is required for the increase in contractile capacity of collecting lymphatics induced by cholesterol depletion with HPßCD, we characterized the contractile function of iaCLVs from Myh11-iCreER^T2^;Cx45^fx/fx^ mice, denoted Cx45^SMC-KO^, under control conditions for 3 minutes and following a single dose with 5mM HPßCD for 60 minutes (Fig. 6A). Interestingly, Cx45-deficient vessels displayed a similar response to cholesterol depletion with HPßCD than that observed in WT vessels (Fig. 6). Despite the loss of Cx45 in LMCs, treatment with HPßCD resulted significant increase in contraction amplitude (Fig. 6A,B) and contraction width (Fig. 6F), and significant decrease in contraction frequency (Fig. 6E). Due to the dysrhythmic, and uncoordinated contractions inherent in Cx45 knockout mice, volume displaced over time was not assessed.

**Figure 6:**
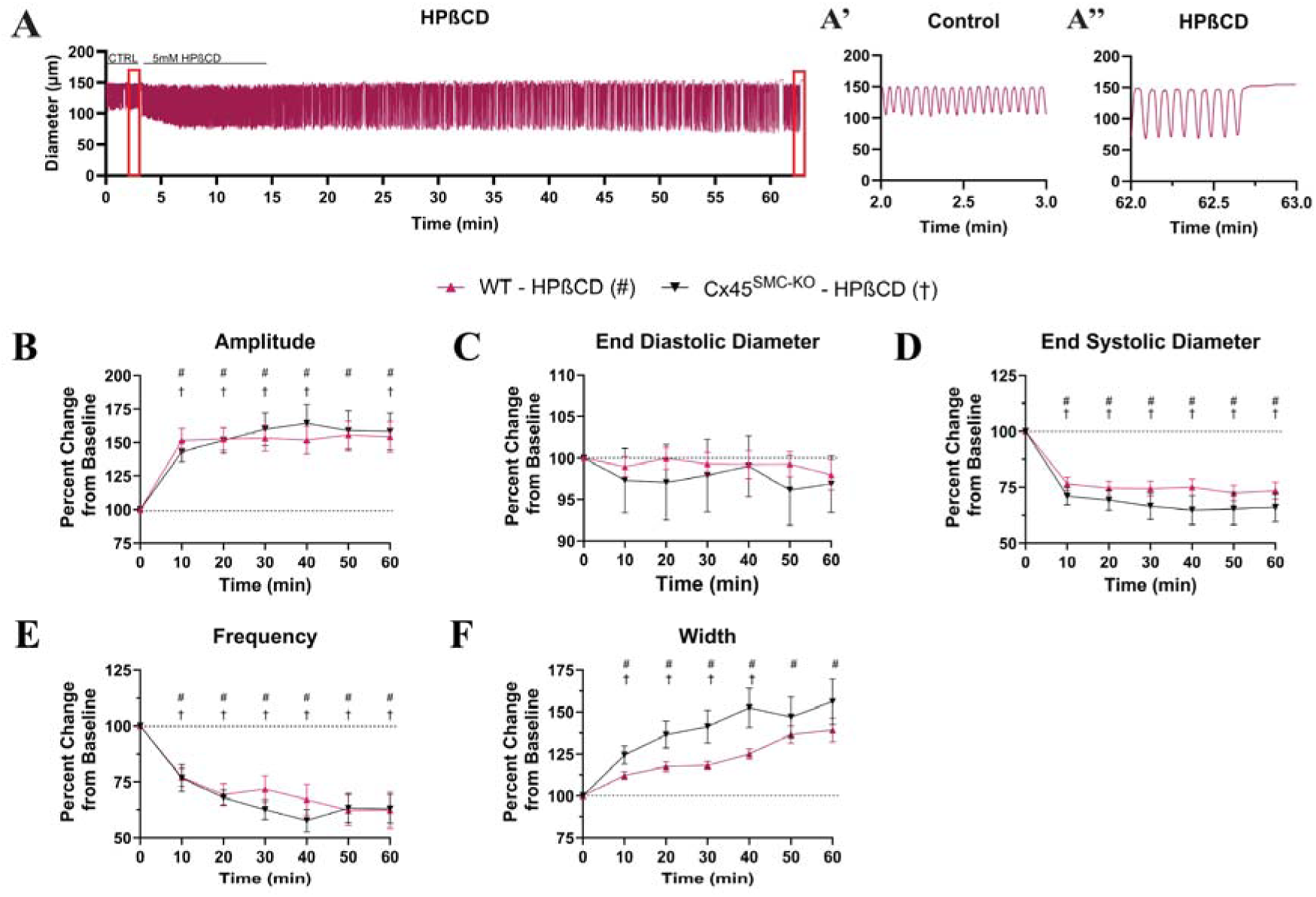
Cyclodextrim-altered contractile function is independent of connexin-mediated cell-cell communication. A) Representative 60-minute diameter tracking of Cx45 KO iaCLV exposed to 5 mM HPßCD treatment. Red boxes highlight 1-minute diameter tracking being expanded in A’) baseline control and A”) after 60 minutes of vehicle or cyclodextrin treatment. Vessel traces were assessed for changes over time for B) amplitude, C) end diastolic diameter, D) end systolic diameter, E) Frequency, F) contraction width, and G) volume displaced. Data are expressed as ±SEM of: WT n=10 animals, Cx45^LMC-KO^ n=5 animals, WT-HPßCD: N=10 vessels, Cx45^LMC-KO^_-_ HPßCD N=5 vessels. A two-way ANOVA with Dunnett’s multiple comparisons test performed. Statistical significance was set at *p<* 0.05 and compared to the baseline for each vessel and are identified as follows: WT-HPßCD (#), Cx45^LMC-KO^_-_ HPßCD (†).

### Increase in membrane cholesterol is detrimental for contractile function

Our results had demonstrated that cholesterol depletion by cyclodextrins led to stronger and more efficient contractions in collecting lymphatics. This would suggest that the opposite, i.e., increase in membrane cholesterol, would be detrimental to lymphatic contractions. To test this hypothesis, we assessed changes to lymphatic function in an acute (cholesterol supplementation by BODIPY-cholesterol) and a chronic (hypercholesterolemic ApoEKO mice) model of increased cholesterol. To acutely increase membrane cholesterol, iaCLVs from WT mice were incubated with 10 µM BODIPY-cholesterol for 1 hour before assessment of contractile function; while for chronic high cholesterol exposure, iaCLVs from 18-20-week-old ApoE^−/-^ knockout mice were used. To reduce animal use, we compared the contractile function of cholesterol supplemented vessels and vessels from ApoE^−/-^ mice to the contractile function of iaCLVs under control conditions shown in Fig. 2. Representative traces of contractions from control, cholesterol supplemented vessels with BODIPY-cholesterol, and vessels from ApoE^−/-^ mice are shown in Fig.7A-C. In contrast to control vessels which displayed large amplitude contractions (i.e., ∼50±17 µm), lymphatics exposed to BODIPY-cholesterol and lymphatics from hypercholesterolemic ApoE^−/-^ mice displayed significantly reduced amplitudes (i.e., ∼20±7 µm and ∼16±6 µm respectively, Fig. 7D). Contraction frequency remained similar in all groups, i.e, ∼17.5±2.4 min^−1^, ∼17.1±3.5 min^−1^, and ∼18.4±2.0 min^−1^ for control, BODIPY-cholesterol, and ApoE^−/-^ respectively (Fig. 7E). The weak amplitude contractions observed in the high cholesterol groups, was also associated with a significant impairment in the volume of fluid that can be displaced by said contractions. Compared to contractions in control vessels, which were capable of displacing a volume of 9.2±3.9 nL per contraction, contractions in BODIPY-cholesterol treated vessels or in vessels from ApoE^−/-^ mice only displaced 3.3±1.0 nL (Fig. 7F). These results indicated that indeed excess cholesterol significantly impairs CLV contractile function.

**Figure 7:**
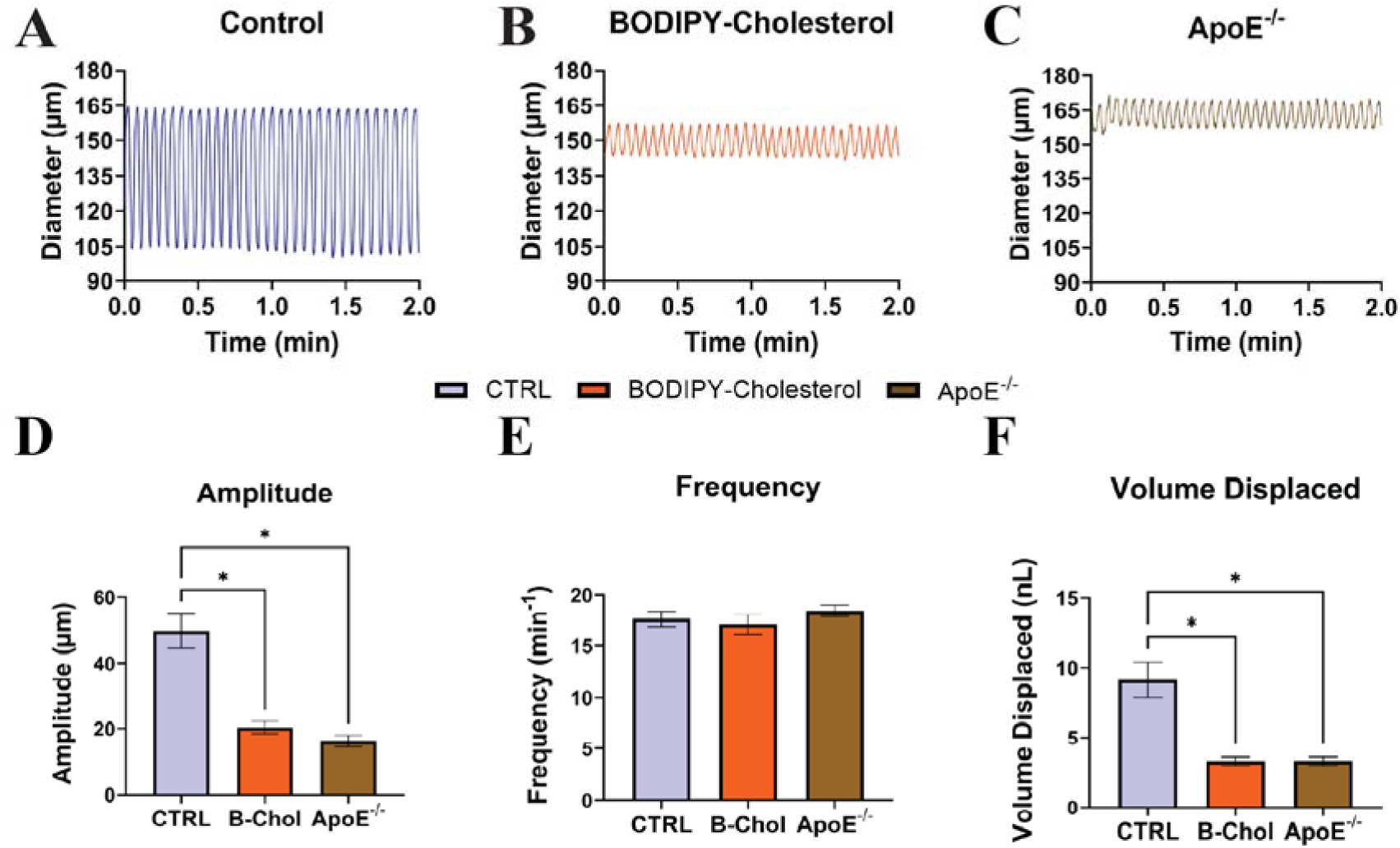
Increased membrane cholesterol impairs lymphatic contractions. Representative trace of a A) WT iaCLV, B) a WT BODIPY-cholesterol-treated iaCLV and C) an iaCLV from an 18-20-week-old ApoE^−/-^ knockout mouse. Traces were assessed for changes in D) amplitude E) frequency and F) volume displaced per contraction. Data are represented as the mean of ± SEM of control: N=10 vessels, BODIPY-Cholesterol: N=12, ApoE^−/-^ N=11 vessels from Control; n=10 mice, BODIPY-Cholesterol: n=5 mice, ApoE^−/-^ n=3 mice with Sidak’s multiple comparisons test was performed. * *p* < 0.05.

### Cholesterol depletion with cyclodextrins ameliorates the high cholesterol-mediated contractile impairment

Finally, we investigated whether cholesterol depletion utilizing HPßCD could restore lymphatic contractile function in both models of acute and chronic hypercholesterolemia included in Fig. 7. Consistent previous sections, we assessed the effects of a single dose stimulation with 5 mM HPßCD to the lymphatic function of iaCLVs supplemented with cholesterol (i.e., pre-treated with 10 µM BODIPY-cholesterol for 1 hour) and iaCLV from ApoE^−/-^ mice. As shown by the representative traces in Fig. 8A,B (including the zoomed-in regions of interest denoted with A’, A”, B’, and B”), the contractile activity of dysfunctional lymphatics from BODIPY-Cholesterol and ApoE-/- groups was significantly improved within the first 30 minutes following treatment with HPßCD. Notably, contraction amplitudes significantly increased by 14.6±2.4 µm in the BODIPY-Cholesterol group, and by 9.1±2.5 µm in the ApoE^−/-^ group (Fig 8C). Interestingly, stimulation with HPßCD resulted in a significant drop in contraction frequency in control vessels from WT mice and vessels from WT mice supplemented with BODIPY-Cholesterol, but it remained unchanged in lymphatics from ApoE^−/-^ mice (Fig. 8C). Treatment with HPßCD led to a significant increase in contraction width and, more importantly, a significant increase in the fluid volume being displaced by each contraction in all groups (Fig. 8D,E). These results confirmed that cholesterol depletion with HPßCD is an alternative to improve lymphatic contractile function in hypercholesterolemia.

**Figure 8:**
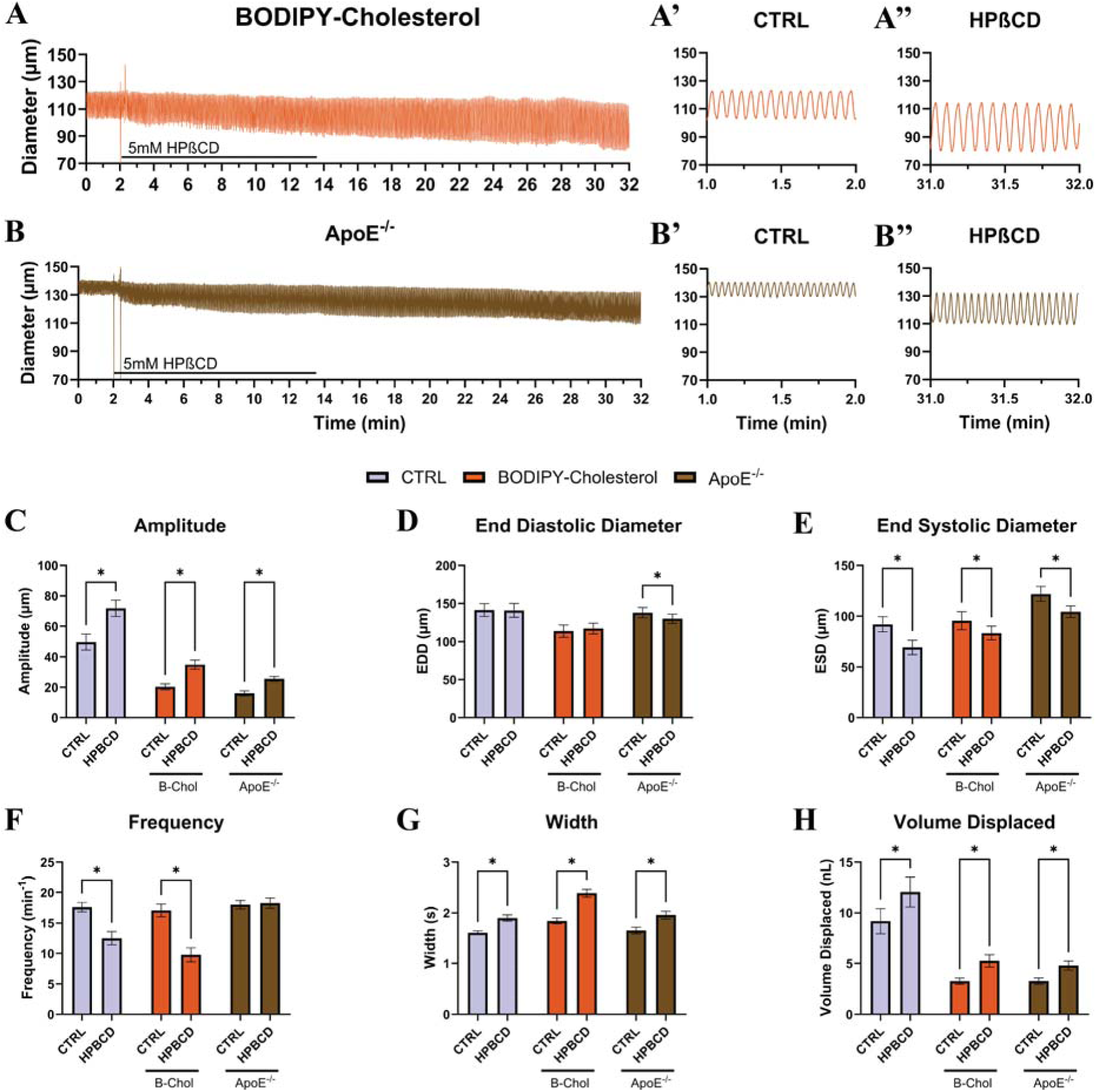
Depletion of excess membrane cholesterol ameliorates impaired lymphatic contractions. Representative trace of A) a WT iaCLV incubated with BODIPY-Cholesterol-treated for 1 hour and B) an iaCLV from an ApoE^−/-^ knockout mouse A’&B’) before and A’’&B”) after cholesterol depletion with 5mM HPBCD. Traces were assessed for changes in C) amplitude, D) end diastolic diameter, E) end systolic diameter, F) Frequency, G) contraction width, and H) volume displaced per contraction 30 minutes after depletion with HPßCD. Data are represented as the mean of ± SEM of control: N=10 vessels, BODIPY-Cholesterol: N=12, ApoE^−/-^ N=11 vessels from Control; n=10 mice, BODIPY-Cholesterol: N=5 mice, ApoE^−/-^ n=3 mice with Sidak’s multiple comparisons test was performed. * *p* < 0.05. with Sidak’s multiple comparisons test was performed. * p < 0.05.

## DISCUSSION

With 20% of the world’s population having hypercholesterolemia^5^ and the presence of excess tissue cholesterol found in lymphedema patients^23^, we wanted to explore the role of cholesterol in lymphatic contractile function and whether this excess cholesterol was involved in the impaired lymphatic contractions seen in animal models of hypercholesterolemia and in lymphedema patients^13,14,23,77^. Here, we established that cholesterol content within the CLVs is involved in maintaining proper lymphatic contractile function and demonstrated the potential role cyclodextrins can play by the improvement of the impaired contractile function in ApoE^−/-^ knockout mice, a model of chronic hypercholesterolemia. To date, there are no FDA-approved pharmacological interventions that significantly improve lymphatic function in chronic diseases such as hypercholesterolemia and obesity-related lymphedema and unfortunately, weight loss alone has proven to be ineffective in correcting the observed dysfunction^78,79^.

Using pressure myography, we demonstrated that cholesterol depletion using cyclodextrins increased CLV contractile amplitude by increasing calcium influx through Ca_v_1.2 channels resulting in a decrease in end systolic diameter and improved in contractile efficiency; however, the decrease in the frequency of contractions at the dose chosen may limit the current therapeutic potential of HPßCD. In vessels with excess cholesterol, ether through BODIPY-cholesterol supplementation or from chronic exposure in ApoE^−/-^ knockout mice, both contractile amplitude and contraction efficiency were impaired while contraction frequency remained unaltered. The impairment to vessel amplitude and contraction efficiency was ameliorated with cholesterol depletion in both acute (BODIPY-cholesterol-treated) and chronic (ApoE^−/-^) vessels, whereas contraction frequency was only impaired in BODIPY-cholesterol-treated vessels, leaving ApoE^−/-^ vessel contraction frequency unaltered with depletion. Together, our data demonstrates cholesterol depletion using HPßCD may be a useful intervention in hypercholesterolemia to restore impaired lymphatic function and the improvements seen in experimental models of lymphedema treated with cyclodextrins^23^ may be due to the rescue of CLV contractile function.

We assessed two different cyclodextrins, MßCD and HPßCD and both of these cyclodextrins are designed specifically for cholesterol depletion. MßCD has been characterized to be more efficient in removing cholesterol and is often the first choice for cholesterol depletion experiments in cell models^55,80^. On the other hand, HPßCD is often described to be more gentle, with a slower depletion rate, and has been approved by the current Food and Drugs Administration (FDA) and the European Medicines Agency (EMA) for phase III clinical trials under the tradename Adrabetadex for the treatment of Niemann-Pick Disease as well as a potential treatment for Alzheimers and Parkinson’s disease^60^. Structurally, cyclodextrins are comprised of a ring of starch with a hydrophilic outer edge and a hydrophobic inner pore^43^. They deplete cholesterol by closely interacting with the lipid membrane, allowing for cholesterol to transfer to the hydrophobic core before they diffuse away. Therefore, the concentration and exposure time of the tissue to cyclodextrins is integral for successful depletion^55^. Studies that utilize cyclodextrins often incubate cells with high concentrations for long periods of time to ensure maximal depletion ^55^; however, longer incubation times and higher concentrations are associated with increased off-target effects such as the depletion of phospholipids and cellular death^55^. To avoid this outcome, we utilized a single washout dose to minimize off-target depletion effects while also remaining in a short therapeutic window and assessed the vessel over the course of the exposure as well as a prolonged period to determine whether the depletion effects resulted in a stable response. Furthermore, this exposure would more likely resemble a response to a subcutaneous injection of cyclodextrin.

In our single dose, washout experiments, within the first 10 minutes of cholesterol depletion, vessels displayed a strong response by an immediate increase in contraction amplitude and a decrease in contraction frequency. These alterations in function implicated a change in the ionic mechanisms that regulate lymphatic contractions. These contractions rely on a slowly depolarizing membrane potential to initiate their intrinsic, phasic response and changes to membrane fluidity through the removal of cholesterol likely altered these ionic mechanisms, resulting in the demonstrated increase in amplitude and decrease in frequency.

Efficient transport of lymph fluid requires the generation of a contraction signal cascade and the subsequent dissemination of that signal to all the LMCs within a vessel segment to efficiently propel lymph forward. This requires the generation of a pacemaker signal and the ability to spread that signal to all nearby cells. LMCs generate their own pacemaker potential^74,81^, however, it is unknown if only a subset of LMCs generate these pacemaker potentials or if all LMCs can generate them. This pacemaker potential is then communicated from the pacemaker site to all surrounding LMCs through cell junctions known as connexins. Specifically, LMCs rely on Cx45 to entrain the contractile wave and produce a synchronous contraction across an entire vessel segment^53^. The loss of Cx45 results in a dyssynchronous contractile wave due to the inability to entrain contractions leading to impaired lymph flow due the activation of multiple pacemaker sites^53^ that propagate slowly through other mechanisms. Importantly, connexins are found within the plasma membrane in cholesterol-rich lipid domains^76^ and interestingly, cholesterol depletion did not uncouple Cx45, as evidenced by the continuous rhythmicity and coordination of lymphatic contractions after depletion. We also wanted to test the hypothesis of whether electrical coupling through Cx45 was required for the improved pumping capacity effect induced by cholesterol depletion with HPßCD; however, there was no difference between the contractile parameters of the WT and Cx45^SMC-KO^ vessels suggesting cell-to-cell coupling through Cx45 was dispensable for the the changes in lymphatic function (increased amplitude and decreased frequency). While cholesterol depletion may not impair established connexin function, changes to lipid rafts could impair the trafficking, assembly, and recycling of new connexins at cell-cell borders. Connexins recycle every 1 to 5 hours and an impairment to the establishment of new connexins would not be observed within our experiments^82^. Long term *in vivo* studies with cholesterol depletion will be required to determine whether lymphatic arrhythmias develop with cyclodextrin treatment. Furthermore, other gap junctions like Cx37 are important for the development, maintenance, and function of lymphatic valves^53,83–85^. Future studies will be required to assess whether cholesterol depletion impairs the permanence and function of the valves.

Altering membrane cholesterol will impact how cells respond to the mechanical stresses applied to them. LECs are known to respond to luminal shear stress and alter lymphatic function through the release of soluble mediators such as nitric oxide and prostaglandins^86–90^, these changes were not assessed in this study and will require further investigation. On the other hand, LMCs respond to circumferential stress generated by changes in luminal pressure^73,91^. We investigated the role cholesterol plays in the LMC myogenic response and found that decreasing luminal pressure resulted in an increased tone response at lower pressures. Myogenic tone is maintained through intracellular calcium levels^91,92^, indicating that cholesterol depletion leads to an increase in calcium within the cytoplasm when experiencing lower luminal pressure after cholesterol depletion and this altered response is lost when pressures return to a more physiological level. Furthermore, the improved amplitude is lost in the lower pressures. This is likely due to the combination of the loss of end diastolic diameter at the lower pressures, impairing their ability to dilate maximally at those pressures as well as the inability to contract further as the end systolic diameter is reaching its lowest maximal diameter, blunting their ability to displace fluid (Fig. 5). Interestingly, intraluminal pressure in mice has been reported to be approximately 4 cmH_2_O^93^, therefore it is unlikely that this impairment of fluid flow at low levels of pressure would lead to impaired lymph transport with *in vivo* treatment.

Cholesterol depletion impairs the sensitivity of many different ion channels leading to an altered ionic response, resulting in the observed decrease in amplitude and end diastolic diameter. Myogenic tone is maintained through a variety of different ion channels including TRP channels, voltage-gated calcium channels, and potassium channels^92^; however, which ion channels are impaired due to cholesterol depletion in CLVs is unknown and a target of investigation. Several studies on cholesterol depletion have identified several candidates^31–33^ such as the TRP channels^94^, Piezo1^95^, and Ano1^96^.

Whether vessels were supplemented with cholesterol by incubating them with BODIPY-Cholesterol or chronically exposed to cholesterol in a model of hypercholesterolemia (ApoE^−/-^ KO mouse model), vessels from both acute and chronic models demonstrated impaired contractile amplitudes and subsequent volume displacement. Of note, contraction frequency is not altered by excess cholesterol in either model, suggesting that the generation of pacemaker potentials is not significantly impaired by hypercholesterolemia nor did vessels display significant contractile arrythmia. Together, the opposing effects cholesterol has on to contractile amplitude implicate the ion channels involved in the generation of the contraction.

Depletion of hypercholesterolemic vessels demonstrated a significant rescue of the impaired contractile amplitude, an increased volume displacement, and a decrease in contraction frequency, however, there was no impairment of contraction frequency in the chronic hypercholesterolemia model (Fig. 8). Together, this suggests cyclodextrin treatment leads to a restoration of some of the ionic mechanisms that were impaired due to cholesterol overload and implicates cyclodextrins as a potential treatment for improving impaired lymph drainage in lymphatic diseases where cholesterol accumulates in tissues such as hypercholesterolemia and lymphedema^13,23^. Further, the improved lymphatic function we demonstrate here likely contributed to the improvement seen in an experimental model of lymphedema treated with cyclodextrins^23^.

Outside of lymphatic contractions, the proper functioning of lymphatic valves is integral to the success of any lymphatic-specific treatment and was not assessed in the current study. It is important to note that the valves in CLVs from mice fed a western diet^77^, valves in ApoE^−/-^ knockout mice^13^ and other lymphatic-altering diseases such as Crohn’s disease^93^ demonstrate significant back leak. Future studies will need to investigate the impact cholesterol depletion has on the lymphatic endothelium and in particular, the lymphatic valves to determine whether they retain their function or even improve their function with cholesterol depletion. Interestingly, the decrease in end diastolic diameter seen in the cholesterol-depleted ApoE^−/-^ vessels indicates a potential positive result. Valve failure resulting in back leak is often due to insufficient valve closure. This can be exacerbated in inflammation by loss of vessel tone preventing the valves from fully closing, and the recovery of some tone may improve valve function and prevent back leak. Improving valve patency will be integral to any lymphatic-specific treatment as proper unidirectional lymph flow requires patent valves. If cholesterol depletion can improve valve patency, the overall function of the lymphatic system would be greatly enhanced despite the loss in contraction frequency and will be explored in greater detail in future work that focuses on LECs.

Here, we demonstrated that the amount of cholesterol within LMC plasma membranes appears to be affecting the function of the ion channels that control contractile amplitude, myogenic response and contraction frequency and having a comprehensive list of LMC expressed ion channels will help narrow down which ion channels may be best investigated, such as voltage-gated calcium channels, TRP channels, other calcium-permeable channels. In fact, the changes we observed were reminiscent of the contractile changes the L-type voltage-gated calcium channel (Ca_v_1.2) agonist BayK8644 induced: an increase in amplitude and a decrease in contraction frequency^35^. Ca_v_1.2 channels are integral to the generation of lymphatic contractions and are responsible for the calcium entry that induces a visible contraction^34,36^. The amount of calcium that enters LMCs through Ca_v_1.2 directly impacts how much of the myosin light chain is phosphorylated and how much it contracts. Increasing the number of Ca_v_1.2 channels recruited or how long the channels are open for will increase the amount of calcium entering the cell, increasing the phosphorylation state of myosin light chain, and induce a stronger contraction. Due to this relationship, we investigated the calcium dynamics within LMCs utilizing the Myh11-iCreER^T2^;Salsa6f mice. Cholesterol depletion not only increased the amplitude but also the duration of each calcium flash, indicating an alteration in calcium handling though Ca_v_1.2. These alterations and the lack of calcium entry with the blockade of these channels implicate them as one of the main channels influenced by cholesterol depletion driving the increase in amplitude.

Investigations into the role of cholesterol in Ca_v_1.2 function have been thoroughly investigated in a variety of different cell types. In skeletal muscle, Ca_v_1.2 channels demonstrate a rightward shift in their activation threshold but have a decrease in L-type calcium current, and suppressed activation and inactivation kinetics^38^. In rat and rabbit ventricular myocytes, cholesterol depletion positively shifted the membrane potential and increased current density, indicating a positive increase in membrane threshold and more calcium flowing through the channel when it was open^41^. In A7r5 arterial smooth muscle cells, cholesterol depletion increased calcium currents, decreased channel inactivation time, and a higher membrane potential was required to inactivate the channeL^97^. Inversely, cholesterol supplementation suppressed calcium currents in coronary macro-arteries, gallbladder smooth muscle cells, and ventricular myocytes^39,41,62,97^, and in A7r5 cells, it did so without altering voltage-dependent inactivation^97^. Together, these studies suggest that membrane cholesterol levels directly impact the function of Ca_v_1.2 channels with low amounts of cholesterol increasing the amount of calcium that flows into the cell as well as increasing when the channels close after they have been open, while inversely, higher levels of cholesterol impair them in a similar but opposite manner.

While Ca_v_1.2 channels are important for changes to amplitude, the loss of contraction frequency with depletion can be influenced by a myriad of other ion channels^74,91^ and past research has implicated that changes to membrane cholesterol can alter their function (reviewed here ^31–33^). Of these channels, several are calcium permeable such as the TRP channels and Piezo1. These small fluctuations of intracellular calcium known as calcium events can alter membrane potential and changes in their function can increase or decrease contractile rate^35^. To determine whether there was a change in the number of these calcium events, we inhibited Ca_v_1.2 channels with nicardipine and counted discrete calcium events in individual LMCs of Myh11-iCreER^T2^;Salsa6f mice (Fig. 4F-H). Interestingly, within one minute of cholesterol depletion, there was a significant decrease in the number of calcium events, which will likely be amplified with longer treatment. While further studies will be required to investigate which type of calcium events are decreasing and which calcium channels are involved, we have demonstrated that cholesterol depletion is modulating calcium dynamics through more than just Ca_v_1.2 channels and together, they can be responsible, at least in part, for the changes to amplitude and frequency described herein.

Outside of calcium channels, many other ion channels can be affected by changes to membrane cholesterol. A few examples of these channels are: Ano1, a calcium-activated chloride channel integral to lymphatic pacemaking^63^ whose activity is increased with cholesterol depletion^96^. TRPV4, an ion channel involved in endothelial regulation of contractile frequency^87^ as well as macrophage release of thromboxane^54^ is augmented with cholesterol depletion and impaired with cholesterol supplementation^94^. Potassium channels such as the ERG channels (KCNH2 or K_v_11.1) that help repolarize the membrane after contraction^98^ also demonstrate an increase in sensitivity to inhibitory drugs with cholesterol depletion^99^. All of which are likely involved in the response to the cholesterol depletion phenotype described here.

While we have focused on investigating the function of the ion channels within the LMCs themselves, we cannot ignore the likelihood that the alterations we see here may be influenced by changes to the LECs. We controlled for the influence of the LECs by conducting our experiments with pressure alone in the absence of flow; however, how the immune cells that are associated with lymphatic vessels respond to cyclodextrins is unknown. Macrophages are found in abundance along CLVs^100–102^ and these immune cells are capable of modulating CLV function through the release of soluble mediators^16,54,103^ and the impact these cells may have on lymphatic function will also require further investigation.

The link between lipids, dyslipidemia, and lymphatic dysfunction is becoming increasingly apparent^12–14,77,104^ and here, we have implicated plasma membrane cholesterol content in LMCs as a primary driver of this dysfunction through impaired calcium entry through Ca_v_1.2 channels. Furthermore, we have identified cyclodextrins like HPßCD as a potential treatment for the reversal of these effects and currently, HPßCD is under investigation as a treatment for a variety of cholesterol handling diseases including lymphedema^23^, atherosclerosis^105–107^, Niemann-Pick type C^60,108^, Alzheimer’s^109,110^, and Parkinson’s^111^. However, the route of administration will be important. Cyclodextrins can be given subcutaneously^112,113^, intravenously^114,115^, or even intrathecally^116^; however, subcutaneous routes will favour the depletion of cholesterol within the local lymphatic vessels first over tissues from more distant sites, allowing for a more lymphatic-targeted approach when given subcutaneously. Depletion of cholesterol depends on the concentration of empty cyclodextrins and the cholesterol content within cell membranes. As the cyclodextrin pores fill, their efficacy in removing cholesterol will diminish. Since lymphatic vessels are unidirectional and drain tissues of excess fluid, a subcutaneous injection of cyclodextrins upstream of the target location would have the greatest impact on the vessels directly downstream from the injection point as these vessels would be exposed to more unoccupied cyclodextrins than more distant tissues such as the aorta or the contralateral limb. However, with impaired lymphatic function, the ability to mount an appropriate immune response is also impaired, will increase the risk of infection from a subcutaneous injection. Studies will be required to determine whether cyclodextrin treatment ameliorates lymphatic function when given from a distant site, especially in humans, but the potential of cyclodextrin treatment in restoring lymphatic dysfunction is promising.

